# A Benchmark of state-of-the-art Deconvolution Methods in Spatial Transcriptomics: Insights from Cardiovascular Disease and Chronic Kidney Disease

**DOI:** 10.1101/2023.12.04.569888

**Authors:** Alban Obel Slabowska, Charles Pyke, Henning Hvid, Leon Eyrich Jessen, Simon Baumgart, Vivek Das

## Abstract

A major challenge in sequencing based spatial transcriptomics (ST) is resolution limitations. Tissue sections are divided into hundreds-to thousands of spots, where each spot invariably contains a mixture of cell types. Methods have been developed to deconvolute the mixed transcriptional signal into its constituents. While ST is becoming essential for drug discovery especially in Cardiometabolic diseases, to date no deconvolution benchmark has been performed on these types of tissues and diseases. However, the three methods Cell2location, RCTD and spatialDWLS have previously been shown to perform well in brain tissue and simulated data. Here, we compare these methods to assess best performance when using human data from Cardiovascular Disease (CVD) data and Chronic Kidney Disease (CKD) from patients at different pathological states, evaluated using expert annotation.

In this benchmark, we found that all three methods performed comparably well in deconvoluting verifiable cell types including smooth muscle cells and macrophages in vascular samples and podocytes in kidney samples. RCTD shows the best performance accuracy scores in CVD samples while Cell2location on average achieved the highest performance across test experiments. While all three methods had similar accuracies Cell2location need less reference data to converge at the expense of higher computational intensity. Finally, we also report that RCTD has the fastest computational time and the simplest workflow requiring fewer computational dependencies. In conclusion, we find that each method has particular advantages, and the optimal choice depends on the use case.

## 1. Introduction

The rise and continued innovation of molecular technologies used within bio-medical research have opened up for novel ways of studying disease and pathogenesis at the cellular level. One such technology is single-cell RNA sequencing (scRNA-seq), which over the past decade has become a central tool for studying cellular heterogeneity at tissue level, thereby gaining crucial mechanistic insights in diseases (Aldridge, 2020). Such insights form the basis for e.g. early phase drug target identification. A more recent technology in the biological research toolbox is spatial transcriptomics (ST). ST, in combination with scRNA-seq, can elucidate how gene expression and specific cell types localize spatially in tissue (Moses, 2022). Understanding cellular migration is key in e.g., inflammation, which is a common disease trait.

A central limitation of multiple ST technologies, including 10X Visium (10x Genomics), is that the resolution is not at the single-cell level, and even for high resolution methods the spatial unit or ’pixel’ is not guaranteed to align with each individual cell. The transcriptomic profile will therefore stem from a mixture of up to approximately 10 cells and often more than one cell type (Figure 1). This challenge has inspired a surge in bioinformatics methods aiming to split up the location-specific transcriptomic profile into its constituents and assign cell types based on reference data, i.e., signal deconvolution. A number of deconvolution methods for ST data have been published (Li B, 2022; Yan L, 2023). In many cases these methods are only validated on healthy brain samples from mice, where cell type populations may be more spatially segregated and well defined (Cable, 2022; Dong, 2021; Kleshchevnikov, 2022). Here we seek to validate methods using cardiorenal disease data, which encompass a spectrum of related disorders of the heart, blood circulation, and kidneys with relation to cardiovascular disease (CVD), which is a complex chronic disease accounting for approximately four million deaths every year in Europe alone, corresponding to 45% of all deaths (Townsend, 2016). ST has the potential to play a major role in unravelling the underlying mechanisms, but currently the described challenges with the resolution level is a major hindrance. Currently, ST plays an important role in target validation and identification. This often requires an accurate connection of gene to cell type where spatial deconvolution is central. Further, deconvolution allows statistical approaches across multiple tissue sections to distinguish artefacts from robust effects in an automated fashion.

**Figure 1:**
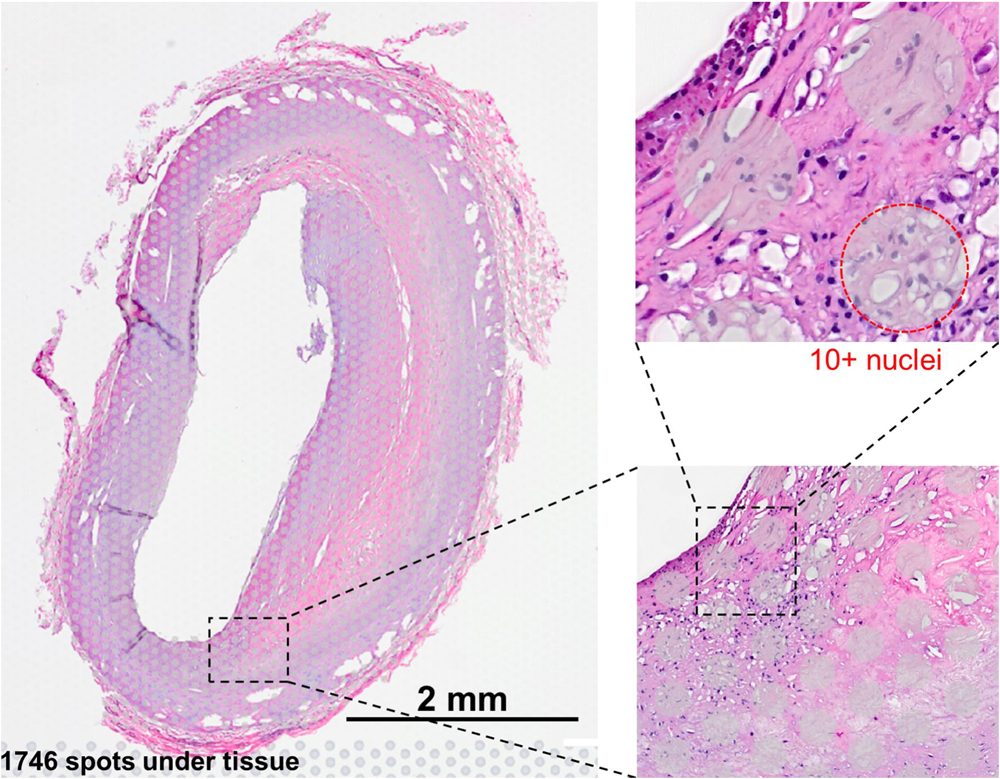
Visium slide showing examples of the barcoded 55um spots where each spot represents a mixture of cells, typically between 5-10 cells.

In this study, we set out to evaluate current state-of-the-art in ST signal deconvolution methods. We applied deconvolution methods to Visium ST data from arterial and kidney samples in healthy and pathological states using three different methods: RCTD (Cable, 2022), Cell2location (Kleshchevnikov, 2022) and spatialDWLS (Dong, 2021), all of which have previously been shown to achieve high accuracy in benchmarks (Li B, 2022; Yan L, 2023). A challenge here is the lack of gold-standards and clinically relevant chronic tissue data. To address this, we performed a benchmark using novel data obtained by manual annotations and subsequently validated the in-silico labels with an expert histopathologist.

## 2. Materials

### 2.1. Arterial plaque and kidney samples

Three publicly available datasets (Alsaigh, 2022; Pan, 2020; Wirka, 2019) were merged to create an atherosclerosis single-cell RNA (scRNA) atlas. The datasets comprised arterial samples from patients undergoing carotid endarterectomy and from heart transplant coronary arteries. The merged atlas consisted of 60,676 cells with nine annotated cell types. It was observed that both RCTD and DWLS experienced memory issues using the full data set when running locally. RCTD has a default setting to downsample reference data on a per cell-type basis, though the choice of sample-size is not validated in depth. To accommodate these issues and improve computational time, an analysis was run to investigate the repeatability of deconvolution results with smaller reference sizes (See results Figure 3.). Going forward random sub-sampling was performed to obtain 3000 cells per cell type. 10 spatial transcriptomics (ST) samples were generated from formalin-fixed and paraffin-embedded (FFPE) samples of coronary arteries, isolated from explanted hearts using the 10X Visium protocol (10X Genomics, 2022/2023). Among the 10 ST samples, five were pathological with clear signs of atherosclerosis, two had early signs of plaque formation, and three were healthy without plaque (Figure 2). For each of the 10 samples, near-adjacent tissue sections (from 1 to 4) from the same coronary artery sample were placed in barcoded capture-areas and processed. The CVD samples ranged from 568 to 1,746 spots, and the CKD samples covered up to 3,966 of the 5,000 total available spots for each capture area.

**Figure 2:**
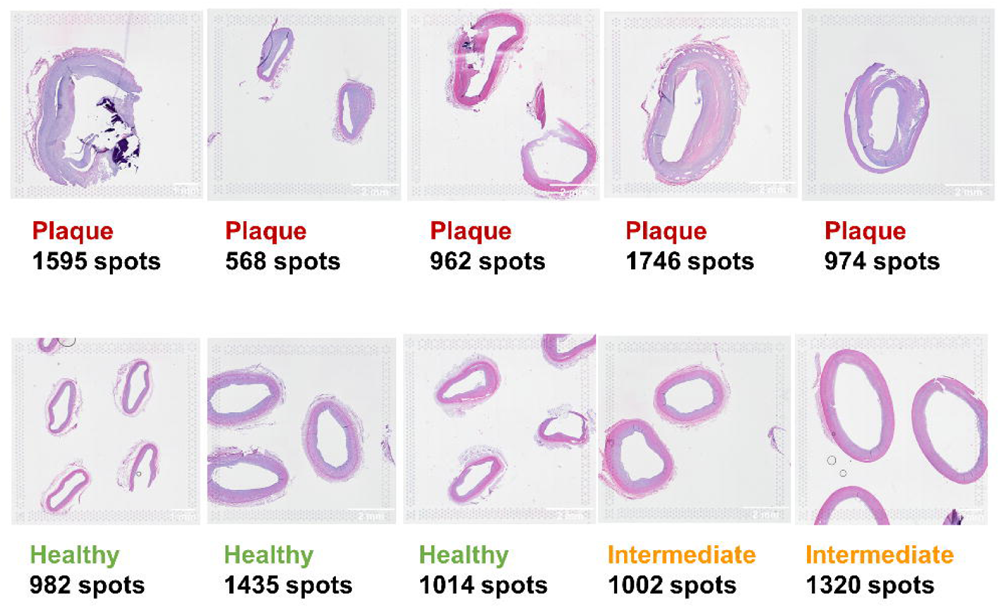
10 Coronary artery FFPE samples analysed with 10X Visium protocol.

**Figure 3:**
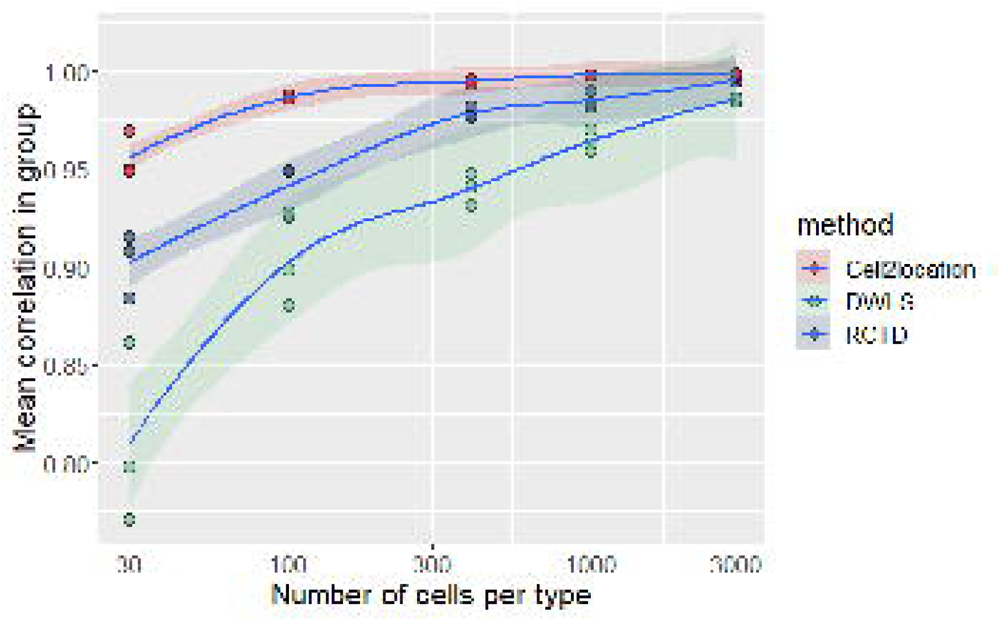
Cell2location deconvolutes sample (CVD2) with higher consistency at a given reference data size.

### 2.2. Visium Protocol

Tissue sections were placed on Visium slides (1-4 serial sections per sample), stained with hematoxylin and eosin (H&E) and imaged using a VS200 slide scanner (Olympus Life Sciences), prior to destainng and overnight hybridization with the Visium human version 1 probe set. The next day, probes that hybridized to mRNA in the tissue sections were eluted and ligated to spatially coded oligonucleotides on the Visium slide. Based on these a cDNA library was created for each sample. Libraries were sequenced on a NovaSeq 6000 (Illumina) sequencing platform, according to the manufacturer’s instructions, using a NovaSeq 6000 S2 Reagent Kit v1.5 (Illumina). Subsequently, reads were aligned with their corresponding probe sequences, mapped to the Visium spot where each probe originally was captured and finally aligned with the original H&E stained image of the tissue section, using the software SpaceRanger version 1.3.0 (10X Genomics).

### 2.3. Benchmarking reference for CVD data

For the benchmarking step, a partial ground truth cell type reference was obtained from an experienced histologist with tissue specific expertise. The ground truth cell type assessment consists of approximately 50 spots per sample, for which it was possible to unambiguously determine a dominant cell type from the available high resolution bright field microscopy images. Across 10 samples, 496 spots were labelled as one of the three cell types: Smooth Muscle Cell (SMC), Macrophage (Mø/MP), or Endothelial Cell (EC). This novel data formed the basis of the benchmark and is as close to a golden standard as possible given the current state of technology.

### 2.4. Benchmarking reference for CKD data

For CKD samples, a scRNA reference was obtained from the publicly available Kidney Precision Medicine Project (KPMP) data repository (Hansen, 2022). Similarly, to the atherosclerosis atlas, random subsampling was performed on a per cell type basis to reduce data set size.

## 3. Methods

We have benchmarked using 3 different models as described below:

### 3.1. Robust Cell Type Decomposition (RCTD)

utilises a Poisson distribution to model counts:

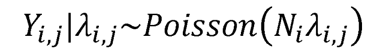

I.e., Given A_i,j_, the random variable Y_i,j_ follows a Poisson distribution with scaling factor N_i_ and rate parameter A_i,j_, where the rate parameter is modelled around the product of the underlying cell type counts and the estimated expression signatures obtained from labelled single cell data.

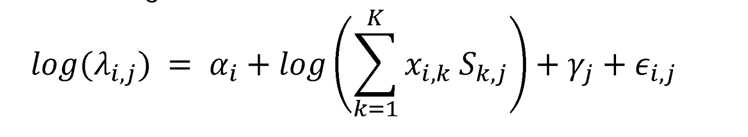

Here a_i_ represents fixed spot specific effects. S_k,j_ is the signature gene matrix, in this case the mean gene expression profile per cell type. The parameter y_j_ represents gene-specific random effects arising from varying gene sensitivity between sequencing technologies. This effect is estimated before other parameters by combining the ST data into a bulk measurement and comparing this to the scRNA expression. E_i,j_ accounts for other random effects or sources of variation including overdispersion.

### 3.2. Cell2 location

is a Bayesian model for deconvoluting ST data into cell type absolute abundances. Cell2location requires two hyper-parameters to be set by the user. The expected number of cells per spot and a regularisation parameter. The first was found by inspecting histology images and counting nuclei for a representative selection of spots and was set to 8 for CVD samples. The regularisation parameter represents the degree to which individual spot sensitivity deviates from the mean within the specific experiment, such that a high value signals consistent detection sensitivity. This parameter was kept at the relatively low default value of 20 to account for varying quality of batches which come from an early exploratory cohort of ST samples.

Cell2location models the observed counts using a negative binomial distribution. The rate parameter is again defined as a function of the signature matrix learned from the single cell reference data and the hidden cell type counts. A spot and gene specific rate are estimated, and parameters are introduced to account for variation in technological sensitivity, shifts due to contaminations, as well as spot specific sensitivity.

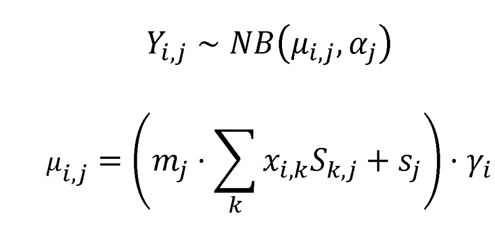

Where μ is the rate parameter and α accounts for overdispersion (equation 3). Parameter mj accounts for technological variation in sensitivity to specific genes. The parameter sj describes an absolute shift in RNA capturing potential gene specific contaminations. Finally scaling parameter γi is included to account for differences in general sensitivity in specific spots (equation 4).

### 3.3. Spatial Dampened Weighted Least Squares,

spatialDWLS, is a Weighted Least Squares approach for deconvoluting cell type proportions of ST spots based on the method DWLS (Tsoucas, 2019) previously published for cell type deconvolution in bulk RNA-seq data.

Similarly, to other methods the observed counts are modelled as a product of estimated expression signatures and the hidden cell type counts of interest.

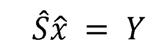

The weighted least squares error minimization problem is defined as:

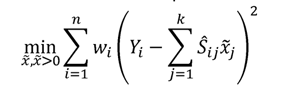

The weights, w, are defined as a function of the dependent variable, the estimated cell-type numbers x, and are therefore determined through iteration with the first iteration being unweighted, Ordinary Least Squares (OLS). This proceeds until the solution converges. The authors introduce a dampening constant to avoid extreme values of weights. This constant is determined by cross-validation of the signature genes to minimise variance.

### 3.4. Influence of reference sample size on repeatability

Sample CVD2 was used for the analyses in combination with different subset sizes of the Atherosclerosis Atlas. Five different subset sizes, with three different replicates of each size, were generated. Random samples were taken without replacement for each of sizes 30, 100, 400, 1,000, and 3,000 cells per cell type. If a cell type had fewer observations available than the chosen sample size, all observations were kept instead. This was relevant for B cells with a total of 2,607 cells, SMC subtypes with a total of 1,016 cells, and mast cells with a total of 764 cells in the atlas. For each of the random subsets, deconvolution of the CVD2 sample was run and the results were compared using Spearman’s correlation coefficient as a measure of similarity of the resulting cell type proportions.

### 3.5. Computational time assessment

For the three methods computational time of training steps and deconvoluting steps were recorded individually. For recording computational time in R, the package tictoc (V.1.1) was used (Izrailev, 2023). In Python the built-in timing functionality of Cell2location’s package was used to record time at each step. The computer used for all computation time recordings was a Windows 10 pc, with a 2.70 GHz, Intel i7-10850H CPU, and 16 GB of RAM. For GPU computation, the NVIDIA Quadro P620, 2GB/512 CUDA core, graphics unit was used.

### 3.6. Benchmarking using histologist provided annotation

The predictive accuracy of each method is estimated by comparing deconvoluted proportions of select spots with annotations provided by a histologist. Cell type annotations available are limited to major anatomically and visually distinct cell populations, this includes smooth muscle cells (SMCs), endothelial cells (ECs) that line the vessel lumen, and in plaque samples aggregated macrophages (MPs). For comparing the histologist-assessed spots (n=496), a dominant cell type had to be determined from the proportions obtained in each deconvolution. Each spot was classified as the cell type with the highest predicted proportion and smooth muscle cell subtypes were grouped. The accuracy (ACC) was calculated per method corresponding to the proportion of true predictions out of all predictions included.

For CKD samples precision and recall was determined and AUROC was calculated using the pROC package (Robin, 2011) based on the histologist annotation of podocyte-containing glomeruli. For a given spot to be classified as containing at least one podocyte a threshold of 15 percent predicted content was used.

## 4. Results

### 4.1. Deconvolution accuracy with variable reference scRNASeq subsets

The effect of reducing reference size on the results of deconvolution was investigated by subsetting of the scRNA data. To assess the confidence with which each method performs deconvolution at a given reference size, a number of replicate analyses were run. In each group of three subsets, the three possible pairwise Pearson correlations were calculated. It was observed that the correlation rose from around 0.8 for spatialDWLS, 0.9 for RCTD and 0.95 for Cell2location, for subsets of 30 cells/cell type, up to at least 0.95 for all three methods at a subset of 3,000 cells/cell type (Figure 3). Correlation was similarly calculated on a per cell type basis. Mast cells, SMC subtypes, and NK-/T-cell estimates generally displayed the lowest correlation within each group, indicating some uncertainty of the results when the reference sample size was too small. All cell types, except Mast cells and SMC subtypes for spatialDWLS, achieved coefficients above 0.90 at the largest sample size. The macrophage/monocyte group was consistently the highest correlating cell type between replicate runs for all three methods (Supp. Figure 1)

### 4.2. Benchmarking method accuracy for major cell types

All three methods performed poorly on observed endothelial cells (ECs). Only DWLS predicted any of the assessed spots as EC dominant, but none in agreement with the ground truth (Table 1). The resulting accuracy scores indicate a similar performance for all three methods. RCTD achieved a slightly higher score than the other two methods at 0.734, compared to 0.702 and 0.708 for Cell2location and DWLS respectively, corresponding to 13-15 additional true predictions. Most endothelial cells were predicted as smooth muscle cells across all methods. Additionally, all three methods predicted a few smooth muscle cells spots as fibroblast dominant, and between 17 and 29 macrophage spots were predicted as smooth muscle cells. Lastly, Cell2location assigned eight macrophages as NK/T-cells.

**Table 1:**
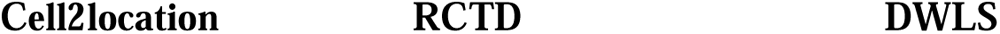
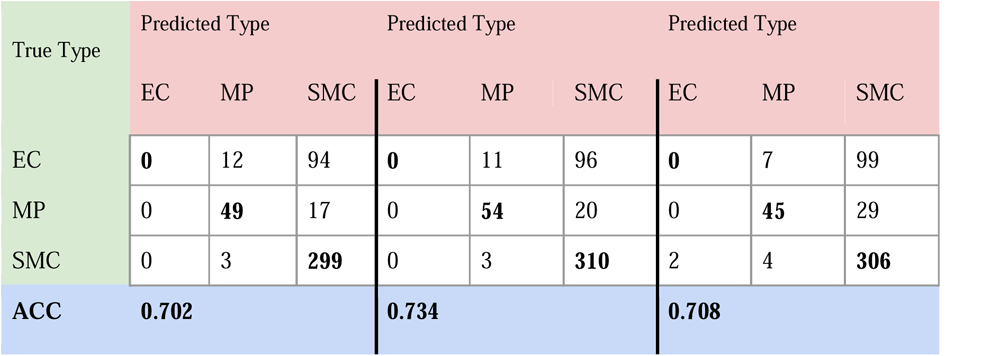
For each method the accuracy was calculated as the rate of success for predicting major cell type in spots with expert supplied ground truth.

| True Type | Predicted Type |  |  | Predicted Type |  |  | Predicted Type |  |  |
| --- | --- | --- | --- | --- | --- | --- | --- | --- | --- |
|  | EC | MP | SMC | EC | MP | SMC | EC | MP | SMC |
| EC | **0** | 12 | 94 | **0** | 11 | 96 | **0** | 7 | 99 |
| MP | 0 | **49** | 17 | 0 | **54** | 20 | 0 | **45** | 29 |
| SMC | 0 | 3 | **299** | 0 | 3 | **310** | 2 | 4 | **306** |
| **ACC** | **0.702** |  |  | **0.734** |  |  | **0.708** |  |  |

### 4.3. Cell type dependent inter-method agreement assessment

Root Mean Square Relative Difference (RMSRD), a metric for pairwise difference between two methods on a per cell type basis was calculated for each pair of methods. A smaller value signifies a smaller mean difference between two methods in relation to the mean proportion of the cell type. A generally greater disagreement is observed for subtype SMCs, NK/T, Mast and B-cells, particularly when compared to DWLS (Figure 4).

**Figure 4:**
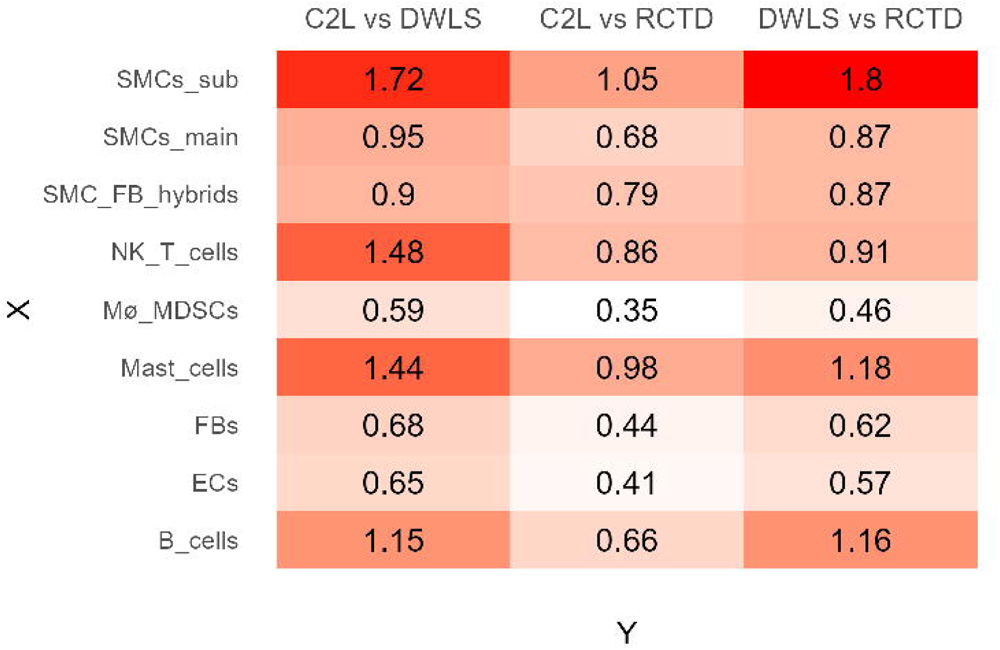
RMSRD between methods for each cell type group. The smaller values in C2L vs RCTD signifies more similar predictions by these methods.

For kidney samples the cell types subject to major variability between methods include Thin Descending Limb (Thin_DL) cells, Parietal Epithelial Cells (PEC), Connecting Tubule (CT) cells, and Ascending Thick/Thin Limb (ATL_TAL). Conversely the rare cell type Podocyte (POD) had high agreement between methods, as did Endothelial Cells (EC), Distal Convoluted Tubule cells (DCT) and Proximal Tubule cells (PT). The previous pattern of higher agreement between Cell2location and RCTD does not seem to be repeated for these samples (Supp. Figure 2).

The high agreement between methods on the podocyte population is of particular interest as it is the most sparse cell type in the reference data (n=244). In comparison PECs are also rare (n=631) but are subject to much higher method-based variation.

### 4.4. Differences in computational time

For each of the three methods, computational times were recorded for a number of samples of varying spot-counts as well as for the different reference subset sizes used. Time spent was recorded individually for any data preparation and then for the deconvolution itself. The data preparation step was defined as all actions that do not need repeating after ST sample is deconvoluted but can be applied directly to the next sample. All three models were found to have a strong linear fit between time spent and the number of spots for deconvolution or number of cells for preparation (Supp. Figure 3).

RCTD and spatialDWLS had similar deconvolution times in the 2-15 minute range for ST samples with up to 3,000 spots. RCTD completed preparations faster, rarely needing more than 5 minutes. Cell2location was much slower spending up to 65 minutes on estimating expression profiles in a 32,000 cell scRNA subset. When deconvolution was carried out using Cell2location, timing ranged from 35 minutes to 2.5 hours per sample.

### 4.5. Benchmarking kidney samples

Given the limited kidney ST samples, it was not possible to perform all identical experiments as for CVD. However, a few experiments were performed to assess the performance of deconvolution. Across three kidney samples, 110 spots were labelled as podocyte-containing glomeruli by expert histologist evaluation of microscopy images as shown in Supp. Table 1. Under the assumption that this evaluation provides a complete ground truth of podocyte locations, performance metrics were calculated across all spots as represented in Table 2. A 10% estimated proportion threshold has been used to classify a spot with a podocyte designation. Cell2location seemed to have higher Precision value as compared to RCTD and DWLS while all 3 methods have similar AUROC values.

**Table 2:** Precision and recall values based on podocyte predictions for each method. As well as AUROC value for podocyte prediction classification threshold using pROC.

|  | Precision | Recall | AUROC |
| --- | --- | --- | --- |
| **Cell2location** | 0.6 | 0.74 | 0.98 |
| **RCTD** | 0.36 | 0.83 | 0.98 |
| DWLS | 0.38 | 0.86 | 0.96 |

## 5. Discussion and Conclusion

The aim of this study was to implement and benchmark a selection of deconvolution methods for internally produced ST data within Novo Nordisk A/S. With a specific focus on human cardiorenal samples at various disease stages, for which limited data has been published. Pipelines were created for the three methods RCTD, Cell2location, and DWLS. Based on scRNA-seq atlas, expression profiles for annotated cell types were estimated following the recommended guidelines for each method. These profiles were then used with their respective method to deconvolute 10 ST samples of arteries and 4 ST samples of kidney tissue at various stages of disease.

In practice, the number of cells necessary to establish a representative cell type profile will vary depending on the heterogeneity of the cell type in question, as well as the quality of the labelling within the reference data. If a cell type is highly homogeneous in the context of the tissue studied, and the instances of this cell type within the reference are accurately labelled, one might expect that a very small number of cells would be sufficient to estimate the expression profile. Conversely, if the cell type is highly heterogeneous, e.g., with multiple unlabelled sub-populations, a larger sample size of representative cells would be preferable to capture as much variance as possible. Despite this, all three models reduce the reference data to a single expression profile for each cell type label, and so for highly diverse cell types, minority sub-populations might be drowned out by the averaging of signals. This is an argument for careful cell type labelling prior to deconvolution, and for dividing cell type labels into subgroups, especially for diverse groups. Also, this illustrates the challenge of approaching a cell type definition as a static expression profile, as clearly, this will be context dependent. Thereby, one well defined cell type may exhibit a multitude of expression profiles.

The variation of cell type estimates from replicate runs were evaluated to validate the choice of subsampling. Most estimates were seen to converge with high uniformity between repeat runs as subsampling neared 1,000-3,000 cells per cell type. Nonetheless Cell2location achieved higher consistency at lower reference data sizes than both RCTD and DWLS, where DWLS struggled with repeatability of specific cell types even at high counts.

It should be noted that since publicly available single cell atlases were used, and we would expect the data to be widely representative for the given tissue. Nonetheless, it might prove beneficial to use reference data generated from the same tissue samples as used for the ST protocol. Therefore, it is also valuable to investigate in future how much single cell or nucleus data is necessary to generate to establish a sufficient reference.

Cell2location and RCTD are both probabilistic methods that rely on discrete probability distributions to model read counts. Cell2location additionally uses a Bayesian probabilistic programming approach. This resulted in remarkably longer computation times observed despite GPU acceleration, but also provided a probabilistic result in the form of a distribution for every estimated cell type proportion, allowing the extraction of cell type abundances at a chosen confidence level. A benchmark was established to compare predictive accuracy. Results indicated that all methods performed well in predicting cell type distributions, at least as far as common cell types were concerned. ECs of the vessel lumen were a problem for all three methods, but it is important to note that the Visium spots are much wider than the endothelial monolayer. A dominant EC signal can therefore not be expected in these spots.

Thus, three methods for deconvolution were successfully applied to internally generated cardiorenal disease data, using the 10X Visium protocol. 10 coronary artery spatial transcriptomics samples were deconvoluted, and all major cell types, including smooth muscle cells, fibroblasts, and macrophages were observed to localize in expected anatomical regions of the arterial vessel. Agreement between methods were found to be high for macrophages, a major cell type of interest in cardiovascular disease.

With ground truth labels supplied by an experienced histology expert, the method accuracies were benchmarked. All three methods achieved similar accuracy with RCTD outperforming the others by a small margin. Cell2location was found to require the least amount of reference data for the supervised step to achieve a consistent output requiring as little as 100 reference cells per cell type for convergent results. Following this analysis, the same workflow was applied to four spatial transcriptomics samples at varying stages of chronic kidney disease. However, due to a smaller number of ST samples available similar inferences could not be made. This could be considered as a limitation of the study given the samples available in the CKD. However, we tried to make sure we could make as much as a sanity assessment in a quantitative manner with the limited CKD samples available. We could observe similar method agreement, and podocytes were identified as a cell type of interest for further analysis. Due to its well-segregated localization in glomeruli only, all three methods offered decent precision metrics of convergence between model and histopathologist expert assessment. Taken together, we think that all three methods are capable of deconvoluting verifiable cell types based on the assessments performed in this benchmarking exercise. It will be interesting to see how these performances will eventually hold or evolve with emerging single cell disease atlases and spatial atlases in cardiorenal areas from public and private consortium initiatives that can also eventually pave a way for disease understanding, drug target discovery and validation. We also encourage the community to perform similar benchmarking initiatives when such large observational cohorts are available at atlas level since this will guide the scientific community not only about the tool’s performance but also where some such spatial technologies are limited.

While it has long been possible to identify cell types of interest by morphology or by targeting a few well-known marker genes or proteins, the newer untargeted ST technologies may be able to provide much higher granularity in cell type and subtype identification. The ability to produce data with near whole-transcriptome coverage and single cell resolution through deconvolution is a very powerful tool that provides a more unbiased and systematic way of studying tissue organisation at different disease states and identifying cell types of interest.

## Data and code availability

Danish legislation regarding sharing of human data requires that the spatial transcriptomics data can be made available upon reasonable request to the corresponding author and approval from The Danish Data Protection Agency.

Scripts for running deconvolutions and output can be found on <u>GitHub</u>.

## Acknowledgements

VD, SB and CP conceived the study. AS prepared the data and ran the Bioinformatics formal analysis. AS prepared the plots and the initial draft of the manuscript under the supervision of VD, SB, CP and LJ. VD and SB performed code review. HH and CP generated the spatial data. CP provided the expert histopathologist assessment. AS, VD and SB interpreted the results. LJ, HH and CP provided additional interpretation of the results. All authors read, edited, and reviewed the final manuscript.

## Supplementary

**Supplementary Figure 1:**
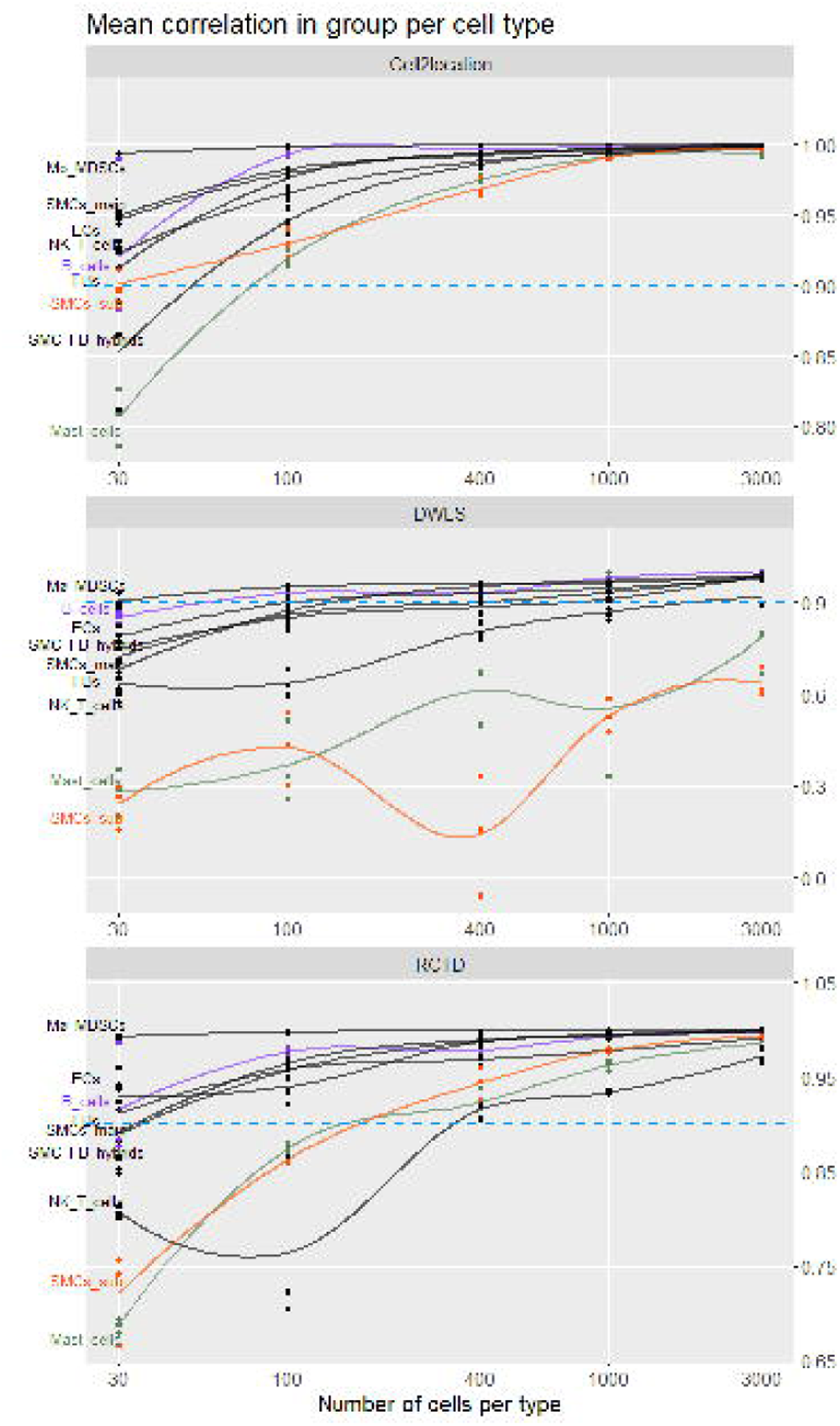
For each method deconvolution of sample CVD2 was repeated with multiple different reference sizes. Correlation between independent deconvolutions was calculated to determine degree of variation in prediction as a function of reference data size. Cell2location is observed to be more consistent across cell types.

**Supplementary Figure 2:**
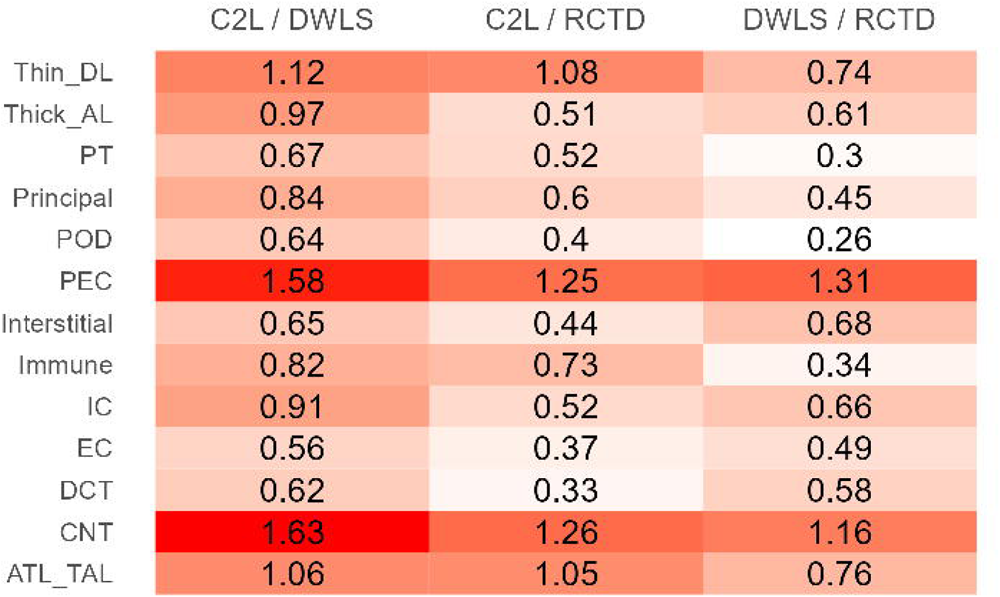
The RMSRD between each pair of methods, with a lower value indication more similar results for the specific cell type.

**Supplementary Figure 3:**
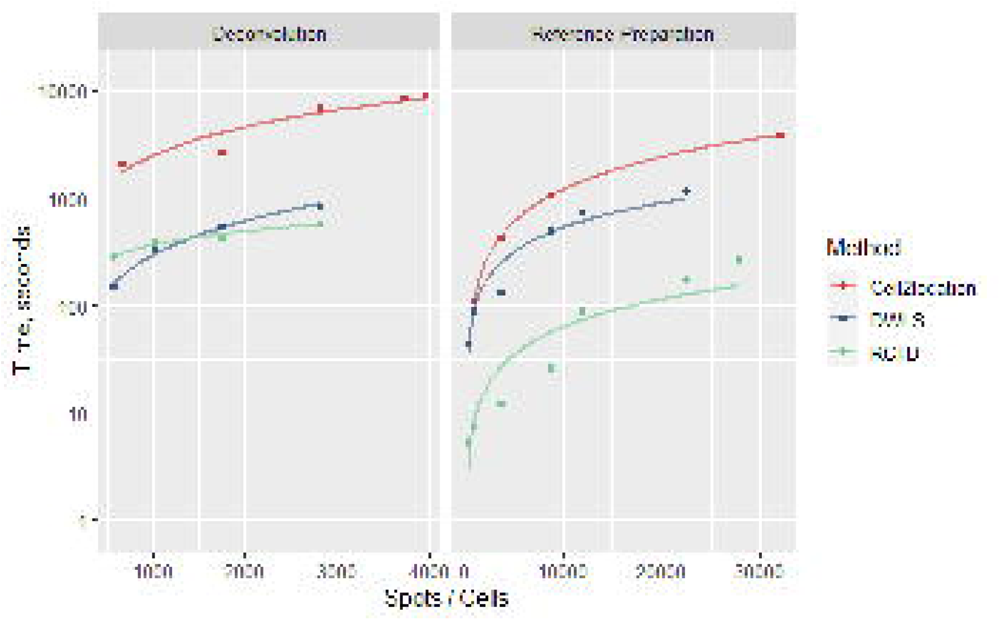
Benchmarking the deconvolution time requirements across methods.

**Supplementary Table 1:** True and false predictions by each method were determined allowing for precision, recall and AUROC metrics to be calculated.

|  | C2L | RCTD | DWLS |
| --- | --- | --- | --- |
| **True Positives** | 81 | 90 | 94 |
| **False Negatives** | 28 | 19 | 15 |
| **False Positives** | 54 | 160 | 152 |
| **True Negatives** | 10997 | 10899 | 10891 |

